# Idea Paper: Predicting culturability of microbes from population dynamics under field conditions

**DOI:** 10.1101/795401

**Authors:** Masayuki Ushio

## Abstract

Isolation and cultivation of microbes from environmental samples have been fundamental and important for species identification and investigating functions and ecology of target microbes. While cultivation and isolation of microbes are not easy, the natural environment can “culture” any endemic microbes, and thus key information for culturing and isolating microbes may be encoded in the natural population dynamics of microbes. In this paper, I present the idea that culturability of microbes may be inferred by quantifying dynamics properties of microbes using nonlinear time series analytical tools. To briefly demonstrate the idea, I analyzed high-frequency, quantitative microbial time series obtained for artificial rice plots established at Kyoto University, Japan. I selected bacterial phyla that included sufficient numbers of microbial taxa, and analyzed 398 microbial taxa using empirical dynamic modeling. The nine phyla analyzed generally followed a similar pattern: many microbial taxa fell into the “Simple” dynamics category, and a small proportion of taxa were categorized in “Simple but nonlinear” or “Nearly random” dynamics categories. The present analysis suggested that many microbes in the study system might be cultivated by modifying a relatively small number of conditions. However, the present idea as well as the result is preliminary and premature, and more precise taxonomic information (i.e., species-level identification) and a culturability dataset will help to validate the idea. If the present idea was found to be valid, *a priori* evaluation of the culturability of microbes would become possible, which would avoid unnecessary costs (labor, time and money) of attempts to cultivate microbes.

## Introduction

Isolation and cultivation of microbial species from environmental samples have been fundamental and important for identifying microbial species and investigating functions and ecology of a target microbial species. However, unfortunately, the majority of microorganisms (e.g., > 99%; note that estimates varied depending on the study) in the environment cannot be cultured easily (e.g., Amann et al. 1995), and improving the recovery of microbes from environmental samples is an important but difficult task. In addition, because cultivation and isolation of microbes are labor-, time- and money-consuming work, *a priori* evaluations of the culturability of microbes would contribute to avoiding unnecessary costs. For example, a cultivation design that includes a large number of combinations of nutrients in media might not be necessary if the target microbes were classified as “easily cultured”.

If *a priori* evaluations of the culturability of microbes become possible, they would contribute to reducing potential costs of cultivation, which would clearly be beneficial for researchers. Also, a framework for quantifying culturability would enable analysis of the relationship between the culturability and genetic factors, which could potentially contribute to understanding why particular microbial species are difficult to culture while others are not.

While isolation and cultivation of microbes are not easy, natural environments can “culture” any endemic microbes, and thus key information for culturing and isolating microbes may be encoded in microbes’ interactions with natural habitats and resultant population dynamics in nature (or under field conditions). Indeed, the diffusion chamber approach, which allows exchanges of chemicals between media and their natural habitat can simulate the natural environment, and improves the recovery of some microorganisms from environmental samples (Kaeberlein et al. 2002, Bollmann et al. 2007). These findings implied that microbes in nature respond to variables in their natural environment, including biotic and abiotic variables, and the interactions between microbes and the environment drive their population dynamics. Therefore, conversely, natural population dynamics of microbes contain integrated information on biotic and abiotic variables, and their interactions, in the microbes’ environment. The question is, however, how we can extract the information encoded in the population dynamics of microbes.

Information encoded in population dynamics (or time series) can be extracted using appropriate time series analytical tools. Recently, modeling tools that are based on attractor reconstruction (Takens 1981) and designed for nonlinear dynamical systems have been developed (e.g., Sugihara et al. 2012), and the framework is called empirical dynamic modeling (EDM). Core tools of EDM are the simplex projection (Sugihara and May 1990) and S-map (Sugihara 1994), which enable quantifying (1) the best embedding dimension and (2) state-dependence of time series data. The best embedding dimension (denoted by *E*) includes information on how many potential variables might be involved in the process (i.e., complexity or dimensionality), and state-dependence (quantified by the nonlinear weighting parameter, *θ*) includes information on how state dependent the process is. A previous study demonstrated that *E* and *θ* are effective indices to distinguish random environmental fluctuations from low dimensional, nonlinear dynamics of organisms (Hsieh et al. 2005). Detailed information on how simplex projection and S-mapping are performed is described in previous studies (Sugihara and May 1990, Sugihara 1994). Also, an overview of EDM is described in Chang et al. (2017).

In the present paper, I present the idea that culturability of microbes may be inferred using population dynamics of microbes and empirical dynamic modeling. Specifically, I expect that the complexity and state-dependency of population dynamics, quantified by *E* and *θ*, would provide information on the culturability of microbes.

## Research hypotheses

I expect three possible dynamics regarding the combinations of *E* and *θ* by following Hsieh et al. (2005): (1) “Simple” (i.e., low dimensional dynamics), (2) “Simple but nonlinear”, and (3) “Nearly random” dynamics (Fig. 1a). First, if the population dynamics of a target species show small *E* and *θ*, then the number of potential variables might be small and the species responds to external forces (e.g., environmental variables and interacting species) relatively linearly (“Simple” dynamics). In this case, the target microbe might be cultivated by examining a relatively small number of cultivation conditions. Second, if the population dynamics show small *E* but large *θ*, then the number of potential variables might be small but the growth of the species is state dependent (“Simple but nonlinear” dynamics). In this case, the target microbe could respond to external conditions nonlinearly, and thus careful considerations of cultivation conditions would be required. Third, if the population dynamics show large *E* but small *θ*, then the dynamics are nearly random and the target microbe might have randomly immigrated from outside the system (“Nearly random” dynamics).

**Figure 1.**
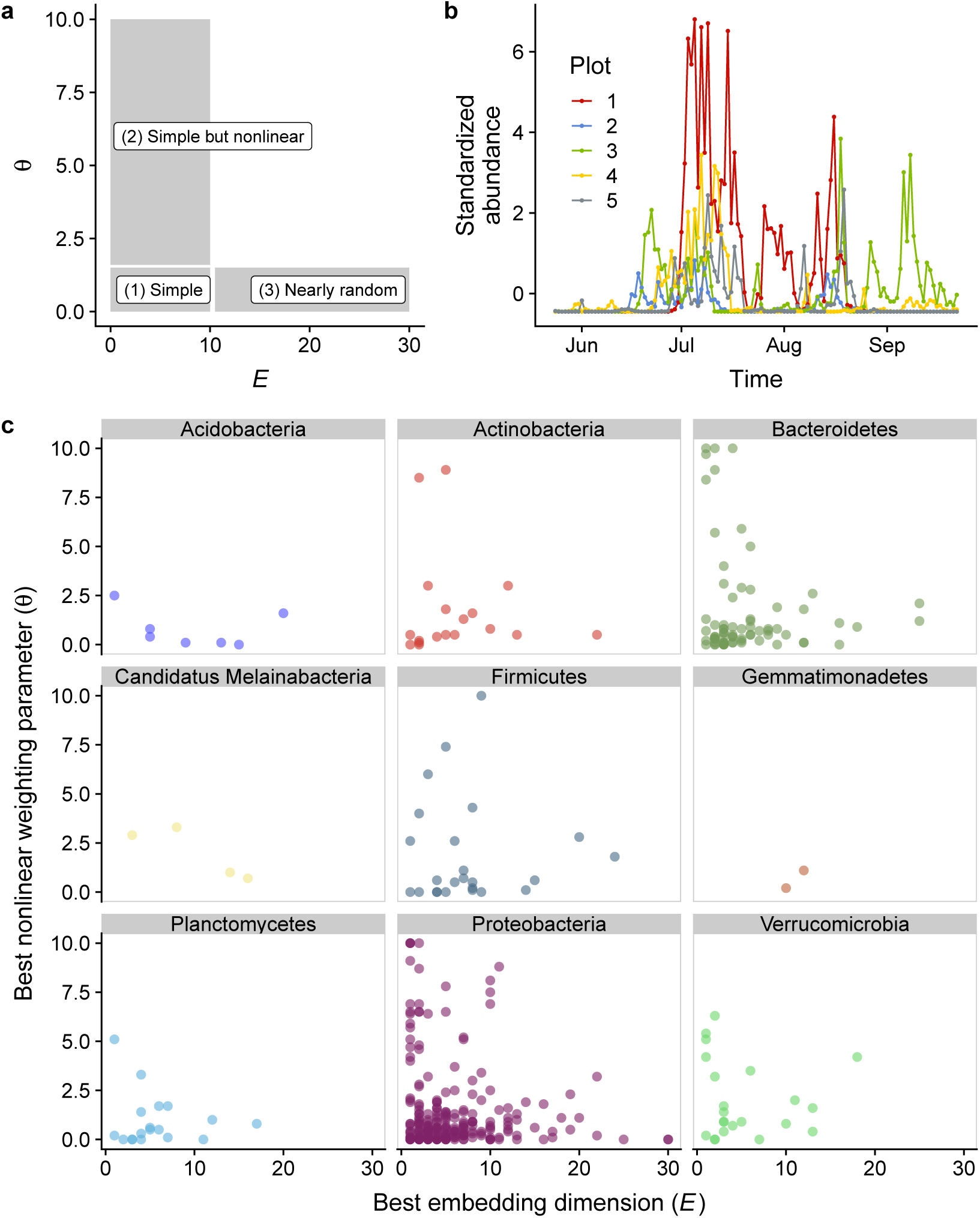
**a** A conceptual explanation of the relationships between E and θ and dynamics properties. Note that the thresholds between the categories are arbitrarily determined. **b** An example of time series analyzed in the present study. Changes in abundance (i.e., estimated DNA copy numbers) of microbial ASVs belonging to Bacteroides are shown. Different colors indicate different rice plots where water samples were taken. c The relationships between E and θ and microbial taxa. Each panel indicates a particular phylum, and each point indicates the best E and θ of each ASV.

## Methods: time series and determinations of the best *E* and *θ*

In order to briefly demonstrate the idea, I analyzed highly frequent, quantitative microbial time series taken from artificial rice plots established in the experimental field in the Center for Ecological Research, Kyoto University, Japan (Ushio, unpublished). Briefly, water samples were collected from five rice plots every day from 23 May to 22 September in 2017 (total 122 days × 5 plots = 610 samples), and filtered using filter cartridges (pore size = *ϕ* 0.22 *µ*m). Then, DNAs were extracted, amplified using 515F-806R primers (Bates et al. 2011) and sequenced using Illumina MiSeq. In the library preparation process, artificial standard DNAs with known concentrations were included, which enabled estimation of the copy numbers of microbial DNAs (“quantitative MiSeq” approach; e.g., Ushio 2019). The sequences were analyzed using the amplicon sequence variant (ASV) approach (Callahan et al. 2016), which has single-level nucleotide resolution and can capture intra-species variations. I selected bacterial phyla that are abundant and contain sufficient numbers of ASVs, which resulted in 398 bacterial ASVs belonging to nine phyla. An example of the microbial time series is shown in Fig. 1b. For the time series selected, simplex projection and S-map were applied to determine the best *E* and *θ* using rEDM packages (Ye et al. 2018) of R (R Core Team 2019). *E* and *θ* that minimize root mean square error (RMSE) of leave-one-out cross validation (LOOCV) were judged as optimal. The whole dataset is being analyzed for different purposes and thus is currently not publicly available.

## Results and discussion

The nine phyla analyzed followed a similar pattern (Fig. 1c): many bacterial ASVs fall into the “Simple” category, and a relatively small proportion of ASVs were categorized as “Simple but non-linear” or “Nearly random”. This result might indicate that, among the selected bacterial ASVs, many microbes could be cultivated by modifying a relatively small number of conditions, which is contrary to the current general consensus that most environmental microbes are difficult to cultivate and isolate (Amann et al. 1995, but see Martiny 2019). This pattern may have been observed because I selected only abundant microbial ASVs detected from the artificial environment. Also, note that undetected, possibly uncultured ASVs were ignored in the analysis because a short-read marker-gene (amplicon) analysis tends to detect cultured representatives more easily than uncultured ones (see debates in the following references: Martiny 2019, Steen et al. 2019).

The present analysis suggested that many microbes in the study system could be cultivated by modifying a relatively small number of conditions. However, the present result as well as this idea is premature and should be verified and developed much more thoroughly. Here I list data that are required to test my idea. First, microbes should be identified at least at the species level. This is because the culturability would be a property of a species, and thus genus or family level identification is insufficient. The present data analyzed were generated by a short-read markergene (amplicon) analysis, and thus the phylogenetic resolution was not sufficient. Recently introduced long-read sequencing would be more appropriate to achieve this goal (e.g., MinION by Oxford Nanopore or Sequel by PacBio). Second, accurate information on which microbial species and how the microbial species is cultivated is required. Information on which microbial species is cultivated would be available in public databases, but developing the cultivation method often requires very detailed tuning and thus the method as well as its complexity would be difficult to fully document/digitalize. Nonetheless, if the present idea were validated, *a priori* evaluation of the culturability of microbes would become possible, which could avoid unnecessary costs (labor, time and money) to cultivate microbes.

## Acknowledgements

I would like to thank Asako Kawai for assistance in the sampling and experiments and Hiroki Yamanaka for providing an opportunity to use Illumina MiSeq. This study was financially supported by PRESTO (JPMJPR16O2) from the Japan Science and Technology Agency (JST) and the Hakubi project of Kyoto University.

